# Cross-platform meta-analysis reveals common matrisome variation associated with tumor genotypes and immunophenotypes in human cancers

**DOI:** 10.1101/353706

**Authors:** Su Bin Lim, Swee Jin Tan, Wan-Teck Lim, Chwee Teck Lim

## Abstract

**Background:** Recent sequencing efforts unveil genomic landscapes of the tumor microenvironment. Yet, little is known about the extent to which matrisome pattern is conserved in progressive tumors across diverse cancer types, and thus its clinical impact remains largely unexplored.

**Findings:** Using a newly generated, unified data resource, we conducted cross-platform assessment of a measure of altered extra-cellular matrix (ECM) composition and remodeling associated with tumor progression, termed as the matrisome index (TMI). Parallel analyses with TCGA in over 30,000 patient-derived biopsies revealed that TMI is closely associated with mutational load, tumor histopathology, and predictive of patient outcomes. We found an enrichment of specific tumor-infiltrating immune cell populations, signatures predictive of immunotherapy resistance, and several immune checkpoints in tumors with high TMI, suggesting potential role of ECM interaction with immunophenotyes and tumor immune escape mechanisms. Both epithelial cancer cells and carcinoma-associated fibroblasts are potential cellular contributors of such deregulated matrisome.

**Conclusions:** Despite wide spectrum of genetic heterogeneity and dynamic nature, matrisome abnormalities are integral to disease progression. Our resource of a curated compendium of 8,386 genome-wide profiles, molecular and clinical associations, and matrisome-tumor genotype-immunophenotype relationships identify potentially actionable immune targets that may guide personalized immunotherapy.

## Introduction

The extracellular matrix (ECM) is a complex multi-spatial meshwork of macromolecules with structural, biochemical and biomechanical cues, influencing virtually all fundamental aspects of cell biology [1]. Although rigidly controlled during normal development, ECM is frequently altered in many diseases, including cancer [2, 3]. Despite clear evidence of abnormal tumor matrix in cancer - owing to complex nature of ECM proteins and their associated factors, or matrisome [4–6] - characterization and understanding their functional role in tumors have been challenging. Little is known about the extent to which matrisome pattern is shared across various carcinomas or unique in tumors of differing metastatic potential. It remains unclear whether there exist subclasses of tumor matrisome that modulates tumor initiation and response to therapy, particularly in the context of immune response.

Previously, we had demonstrated a robust association between a pattern of 29 matrisome genes and its predictive value in prognosis and adjuvant therapy response in early-stage nonsmall cell lung cancer (NSCLC) [7]. This specific pattern of matrisome gene expression is termed the tumor matrisome index (TMI). Given that abnormal ECM dynamics are a hallmark of cancer [3], we hypothesized that this pattern of genomic aberrations in tumor matrix is associated with intrinsic resistance to the therapy, patient outcomes and other clinically relevant parameters, across various tumor types. The present study thus extended evaluation of the TMI to 11 major cancer types (Fig. 1A).

**Fig. 1.**
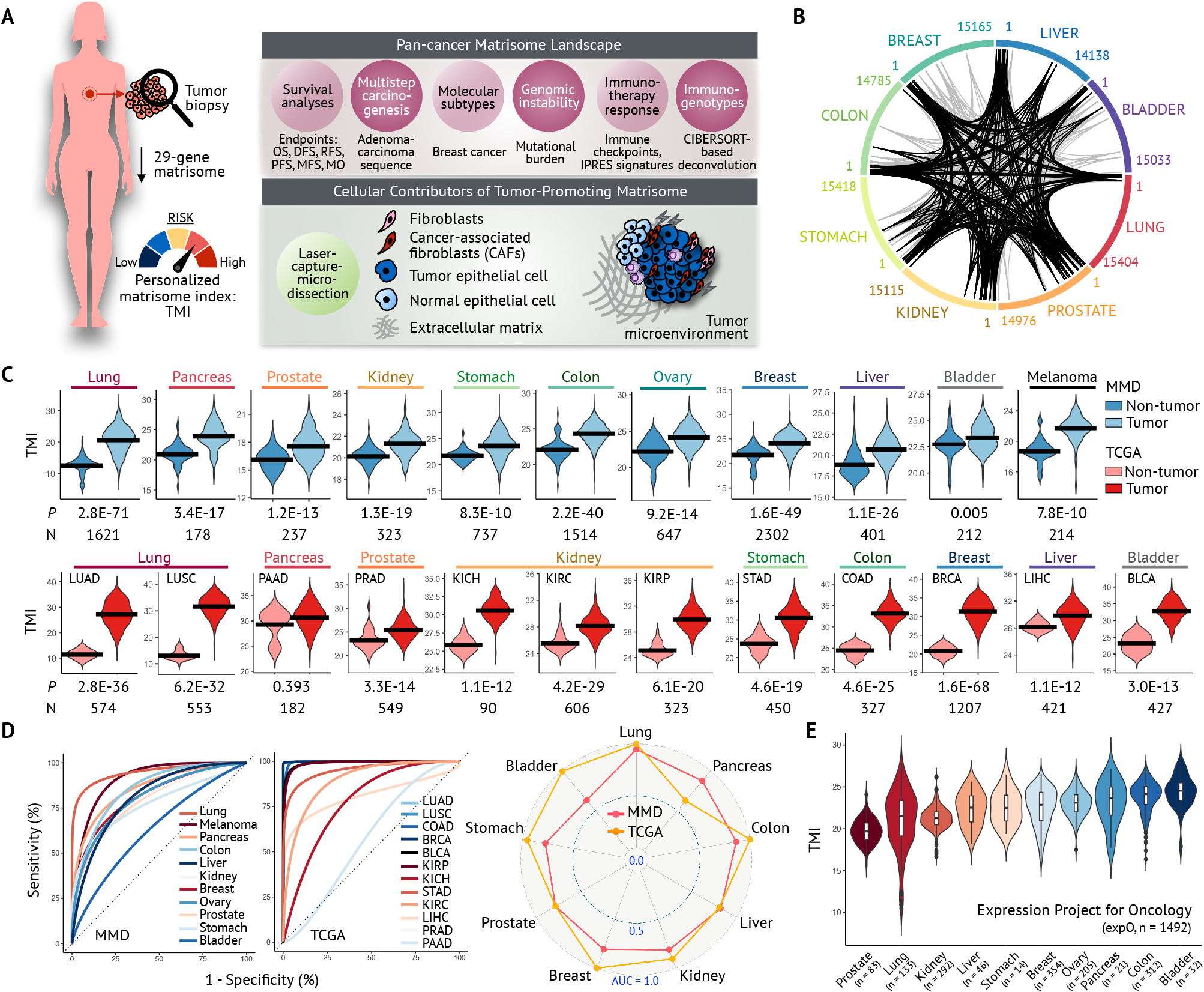
Conserved pattern of deregulated matrisome expression in various carcinomas. (A) Study design: cross-platform pan-cancer analysis of TMI across 11 cancer types using microarray-acquired-MMDs (*n* = 8,386) and RNA-seq-acquired TCGA (*n* = 5,709). (B) Circular plot illustrating the ranked position of matrisome genes based on differential expression (cancer vs. normal) using TCGA (1 being the most differentially expressed). Black and grey lines show generic and lung-specific matrisome signature, respectively. (C) Comparison of individual TMI in tumor and tumor-free tissues. *P* = the Mann-Whitney U-test *P*-value; N = number of samples analyzed. (D) Diagnostic accuracy of TMI; smooth ROC curves (left) and AUC are shown (right). (E) Inter-tumor variation of matrix remodeling across 11 cancer types using an independent data source (see Methods).

To facilitate cross-platform meta-analysis of TMI with TCGA, we generated a single, cancer type-specific, merged microarray-acquired dataset (MMD), comprising thousands of transcriptome profiles of both tumor and tumor-free tissues (Tables S1 and S2). Given the derivation from a uniform computational pipeline, MMD accounts for technical bias from preprocessing.

We further define the pan-cancer matrisome landscape by exploring previously unknown features of matrix remodeling and their impact on tumor genotypes and immunophenotypes. Using this newly curated MMD together with 12 TCGA cohorts, we found robust associations of tumor-promoting matrix composition with genomic instability, molecular and clinical features, and immune infiltrate composition. At the center of our results is the promising clinical applicability of TMI to predict immunotherapeutic responsiveness and the identification of specific immune targets that may be selectively efficacious in subset of matrisome-deregulated carcinomas.

## Results

### Patterns of deregulated matrisome genes are conserved in various malignancies

Given that the initial TMI was derived from lung cancer, we first questioned the extent to which genome-wide DE patterns of lung cancer would be shared across various carcinomas. Conducting cross-platform analysis with TCGA, we found that DE patterns of lung cancers were comparable to that of breast, ovarian, bladder, colorectal, and prostate cancers, regardless of assayed platforms (Tables S3 and S4). Recent pan-cancer studies have revealed some cancers of these tumor types were classified into common molecular subtypes [8, 9]. Assessing the initial 29-matrisome-gene signature at the individual gene level, we found that many genes were consistently positioned in the top of the ranked DE gene list across multiple epithelial cancers, despite their initial derivation from lung cancer (Fig. 1B and Table S5). We further identified a subset of these genes, or “generic” signature, that was significantly enriched in most cancers (Table S6; see Methods). This initial screening provided support for a valid hypothesis that altered matrisome dynamics is a common tumor response across various cancer types. We thus investigated whether previously developed scoring metrics (TMI; see Methods) could further serve as a clinical tool for predictive medicine.

### TMI classifies cancer

Analyzing a total of 14,095 biopsies (Tables S7 and S8; see Methods), all eleven cancers examined demonstrated significant difference in the TMI between tumors and tumor-free tissues, except for TCGA PAAD (Fig. 1C). Non-significance in PAAD is likely due to insufficient normal samples for comparison (normal n = 4 vs. tumor n = 178). Lung cancer-derived TMI expectedly achieved almost perfect accuracy in lung cancer diagnosis (MMD AUC = 0.946 and TCGA AUC = 0.999; Fig. 1D and Table S9; see Methods). Robust diagnostic potential was also demonstrated across all tumor types at any given time and tumor purity, to different extents. The observed difference in some cancer types between two assayed platforms is due to different number of genes filtered in TCGA for final index computation (Table S2).

A further independent data source where tumors were procured under standard conditions and examined 1,492 tumor samples across 10 cancer types was examined (Fig. 1E and Table S10; see Methods). Although over a narrower range than for lung cancer, the rest of carcinomas displayed a broad range of the index, revealing a degree of heterogeneity and spectrum of matrix remodeling at different stages of cancer progression. Given minimal technical variation introduced between samples from this single resource, the consistent observed difference in the range of index and wide biological variability suggest a characteristic and causal mechanism and role of ECM molecules independent of site of tumor origin.

### TMI is predictive of patient outcomes

Using 72 independent datasets, we found that the predictive power of TMI for patient survival was dependent on cancer type (Table S11; and for lung cancer [7]). The TMI was an unfavorable prognostic factor for overall survival (OS) across prostate, melanoma, liver, pancreatic, renal, and breast cancers (Fig. 2A-B, Figures S1 and S2). Although not consistently significant, predicted low-risk patients with colorectal and bladder cancer had better survival for a number of patient cohorts. In contrast, there was no negative correlation between the index and survival in patients with ovarian and gastric cancer; rather TMI was a positive prognostic factor for recurrence and survival in gastric cancers.

**Fig. 2.**
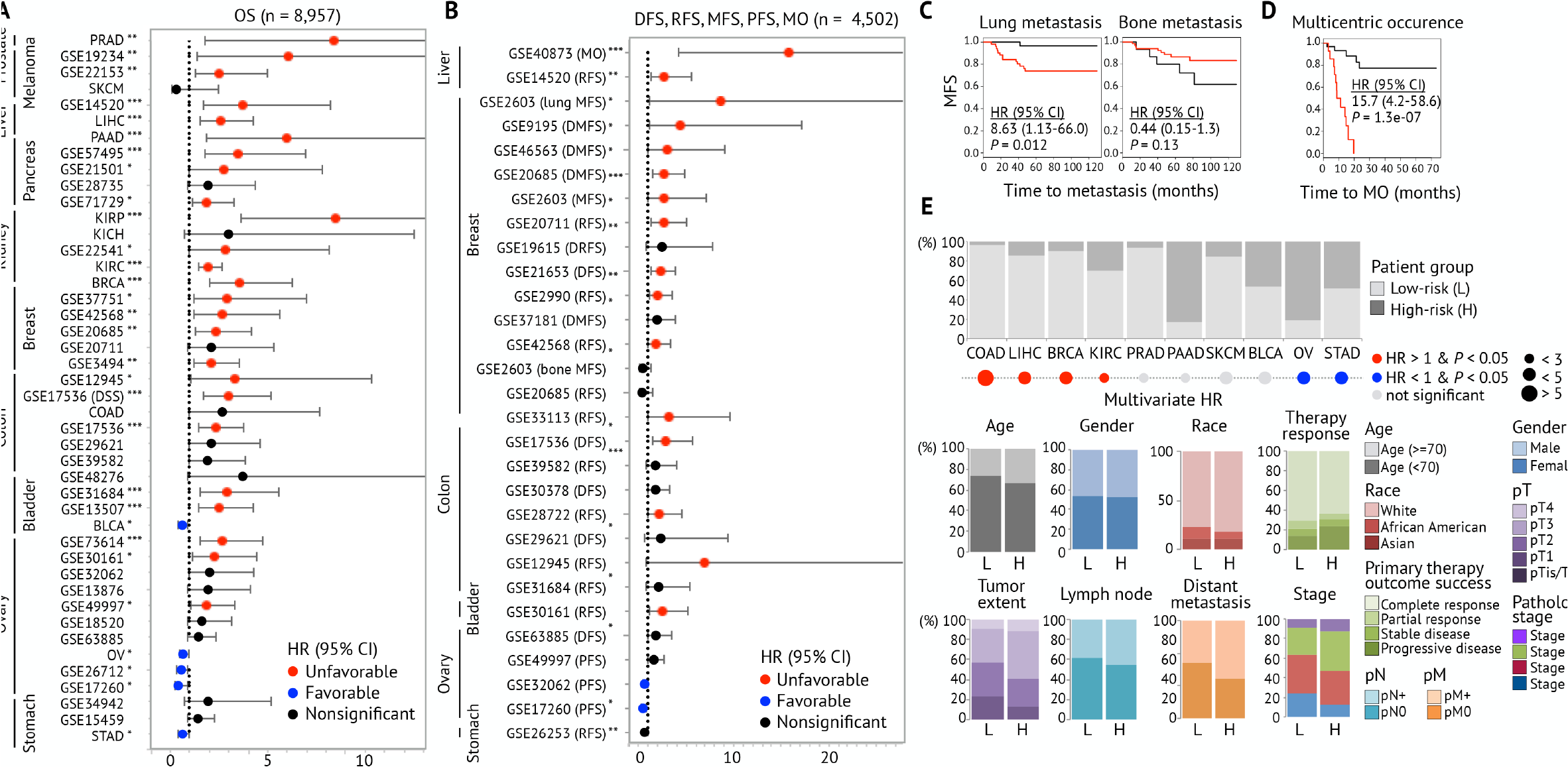
Clinical features associated with TMI. (A) HR forest plot for OS and DSS. (B) HR forest plot for other surrogate endpoints (abbreviated names are provided in Table S12). *P* = log-rank *P*-value; *n* = number of samples analyzed. (C) Lung- and bone-metastasis-free survival curves of breast cancer patients (GSE2603, *n* = 82). (D) Multicentric occurrence-free survival curves using noncancerous tissues of early-stage hepatocellular cancer patients (GSE40873, n = 49). Univariate HR with 95% CI and log-rank *P*-value (*P*) are stated. Black and red KM curves represent stratified patients in low-and high-risk group, respectively. (E) Multivariate Cox regression analyses using TCGA and comparison with conventional clinical parameters between two groups stratified based on the TMI (Table S13).

Clinical associations with not only OS but also progression-free survival (PFS) and metastasis-free survival (MFS) provides better insights into the matrisomal changes during tumor progression and metastatic dissemination. For example, the index was highly correlated with metastasis development in lung (HR = 8.63, P = 0.012), but not in bone (HR = 0.44, P = 0.13), among breast cancer patients (Fig. 2C); this is consistent with the proposed role of extracellular protein encoding genes, such as proteases and chemokines, in promoting metastatic potential [10]. Liver cancer, among others, demonstrated marked predictive capacity of the index for multiple surrogate endpoints (Fig. 2D). Its potential use as a predictor of multicentric occurrence for early-stage hepatocellular carcinomas may guide clinical course and treatment strategies, which depend on classification of subtype of multifocal HCC tumors [11].

Multivariate analyses taking into account age, race, gender, and clinical and pathological TNM status revealed that TMI is an independent predictor of survival in colorectal, liver, breast, renal, ovarian, and gastric carcinomas (Fig. 2E and Table S12). Comparing with traditional clinical features between two stratified patient groups in TCGA, we found that the high-risk group had a higher proportion of patients staged pathologically as T3 or T4, diagnosed as lymph node and distant metastasis positive, and classified as late stage.

### There exists a tumor genotype-matrisome phenotype relationship

Given the above association with disease progression and stage of cancer, we asked if there was progressive dysregulation of the TMI in the step-wise progression to advanced cancer. Expression of progression markers should gradually increase or decrease during multistep carcinogenesis to be clinically applicable [12]. Across breast, colorectal, and pancreatic cancer, we found a progressive increase in the index during tumor development from normal to adenoma to carcinoma in-situ, and further to invasive carcinoma (Fig. 3A and Table S13).

**Fig. 3.**
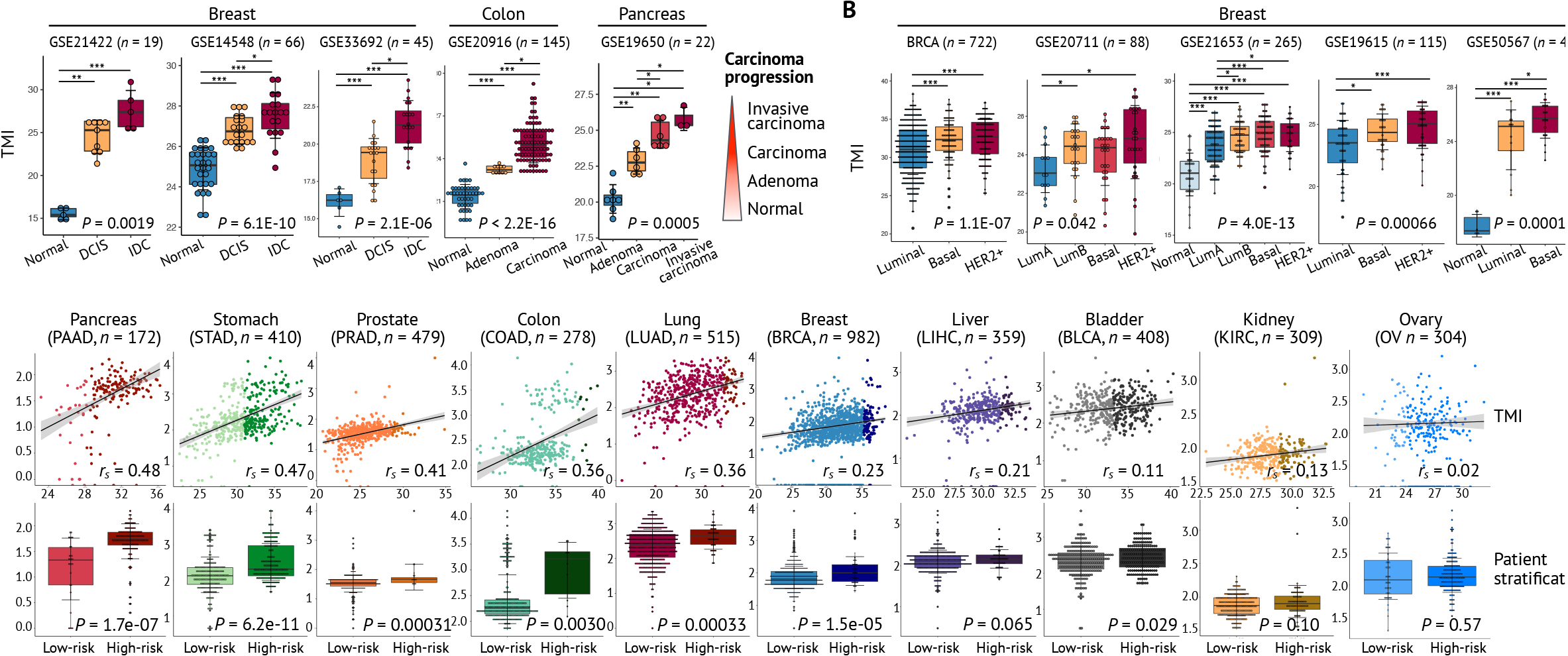
Histpathological and molecular features associated with TMI. (A) Change in TMI during multistep carcinogenesis. (B) Association with molecular subtypes of breast cancers: normal breast-like, luminal A and B, basal-like, HER2-enriched. (C) Correlation with mutational load using TCGA. Patients are stratified into low- or high-risk group based on the TMI. Linear regression lines are drawn (black line) with 95% CI (grey zone). \*\*\**P* < 0.001, \*\**P* < 0.01, \**P* < 0.05 using the Mann-Whitney U-test comparing two groups; Kruskal-Wallis *P* values are stated; *n* = number of samples analyzed.

Molecular subtyping in breast cancer is often associated with its predictive value for prognosis and response to various treatment options [13]. We found that basal-like and HER2+ tumors consistently had higher TMI than luminal tumors (Fig. 3B and Table S14); paralleling trends were reported in prior works for breast cancer specific survival [14, 15]. Collectively, breast cancers exhibit the strongest association of TMI with both molecular and clinical features among all types of cancer investigated.

Using TCGA datasets, we found a strong correlation of the TMI with tumor mutational burden (TMB), defined as the total number of simple somatic mutations, in pancreas, stomach, prostate, colon, lung, and breast cancers (Fig. 3C, Tables S15 and S16; see Methods). This suggests a potential role of matrisome molecules in providing unique selective pressures in the tumor microenvironment and increasing genomic instability. Given that non-synonymous somatic mutations are major determinants of tumor immunogenicity in many solid tumors [16], we next asked if there exists any direct association of this abnormal matrisome with the immune landscapes and conducted cross-platform evaluation of TMI in the context of immune response.

### Matrisome associates with tumor immunophenotypes and escape mechanisms

By applying CIBERSORT to MMDs and TCGA, we found that for many cancers, TMI was closely correlated with composition of specific tumor-infiltrating lymphocyte populations (Fig. 4A). Enrichment of these TILs related to both innate and adaptive immunity was diverse and cancer-specific. Relative abundance of M0 and M1 macrophages, neutrophils, activated mast cells, regulatory T cells (Treg), and T follicular helper (Tfh) cells, activated CD4+ memory T cells generally increased, whereas that of resting CD4+ memory T cells, mast cells, naïve B cells, and resting dendritic cells decreased along with tumor-promoting matrix remodeling. Impact on immune infiltrates was particularly pronounced in gastric, lung, colorectal and breast cancers, having significant positive and negative correlations with TMI. This highlights the evolving nature of the immune response during matrisome remodeling in these selected cancers. It is noteworthy that despite having no correlation with mutational load, TMI of some cancer types, such as liver, renal, and ovarian cancers, remains associated with specific TIL composition. This indicates that not only tumor genotypes but also matrisome phenotypes may be critical determinants of tumor immunogenicity and thus inter-tumor heterogeneity of immune infiltration, considering highly heterogeneous expression of ECM molecules [7].

**Fig. 4.**
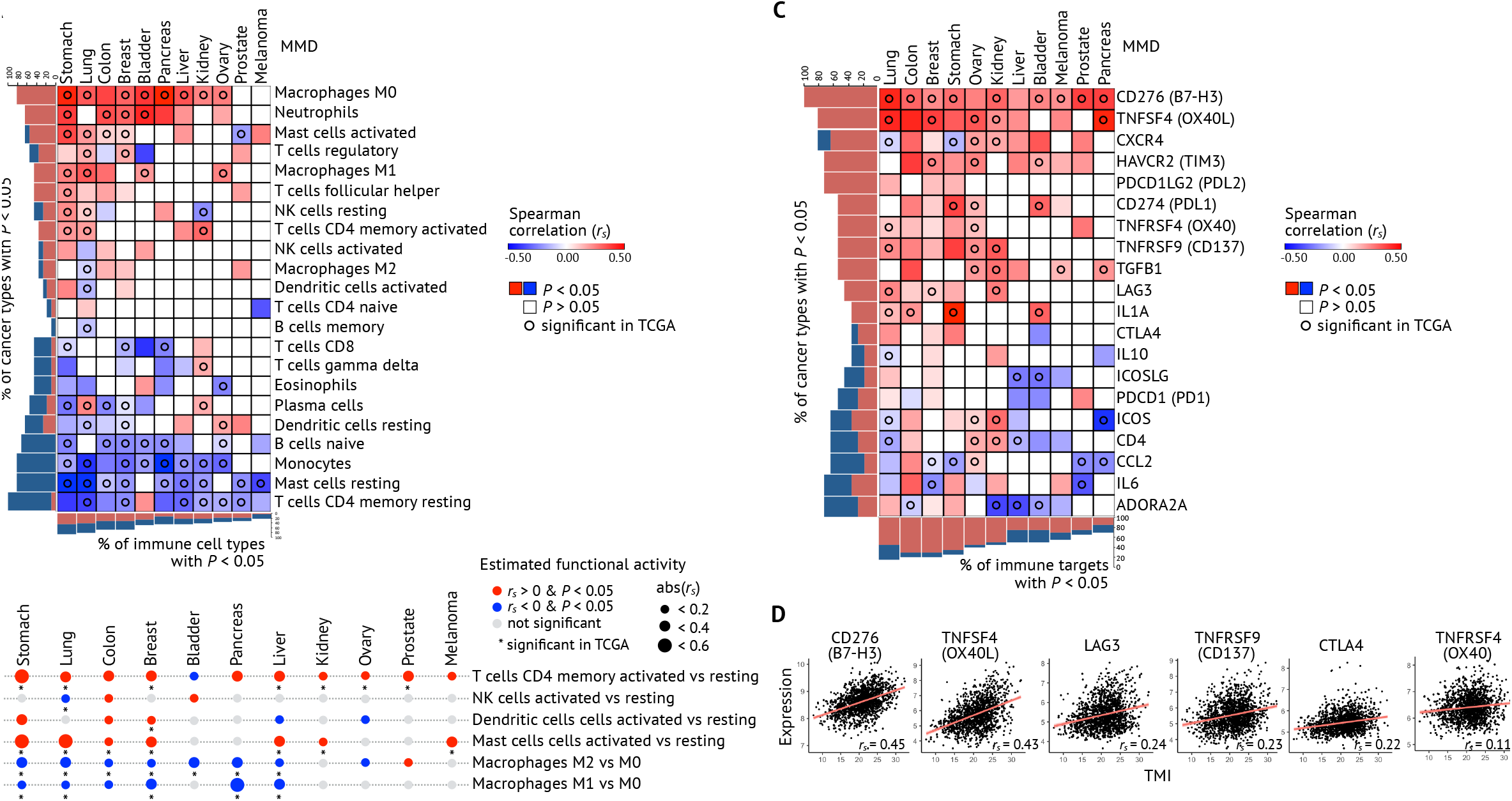
Cross-platform evaluation of TMI in the context of immune response. (A)Correlation between TMI and CIBERSORT-estimated relative abundance, and (B) functional activity of 22 immune cell types. (C) Correlation between TMI and expression of 20 clinically targetable immune checkpoints; *r*_*s*_ = Spearman’s correlation coefficient. (D) Linear regression lines (black line) with 95% CI (grey zone) are drawn for selected immune targets with statistical significance (*P* < 0.05) in lung cancer. *r_s_ =*Spearman’s correlation coefficient.

We further estimated functional activation of distinct immune cells, as recently described [17], for their potential role in specific killing of tumor cells upon immunotherapy. Across all cancers except for bladder cancers, the activation state of CD4+ T cells was significantly higher in tumors with high TMI, suggesting their central role in response to abnormalities in the local matrix niche (Fig. 4B). Activations of other innate immune infiltrates such as NK cells, DC cells, and mast cells, were also either positively or negatively correlated in specific cancer types, whereas in others these cell types had no correlation with matrisomal changes. Due to high enrichment of M0 macrophages, functional activities of M1 and M2 macrophages were repressed in tumors with high TMI (and thus classified as high-risk based on cut-off index), shifting the tumor microenvironment toward an immunosuppressive phenotype.

We next correlated the index directly with the expression of 20 potentially targetable immune checkpoints [18] – that are currently in preclinical or clinical trials, and are FDA-approved. Many of these genes were enriched in tumors with high TMI, potentially responding to expanded CD4+ T cells (Fig. 4C and Table S17). The highest number of immune targets with correlation was found in lung cancer; those genes whose expression levels are positively correlated with TMI in both MMD and TCGA are shown in Figure 4D. Particularly, B7-H3 is a promising pan-cancer immune marker, which may be selectively efficacious in these high-risk tumors. Altogether, these findings unveil specific immune targets that may be selectively efficacious in a subset of matrisome-deregulated carcinomas. We expect that these tumors will be more sensitive to immune checkpoint blockade given that they harbor sufficient immune infiltrates and upregulated immune checkpoints.

Given promising applicability of our metrics in prediction of immunotherapy responsiveness, we tested how this specific matrisome would be expressed relative to a previously defined signature of cancers that fail to respond to immune checkpoint inhibitors. By applying innate PD-1 resistance signature (IPRES) [19] to lung MMD and TCGA LUAD, we found that patients who are predicted to be resistant, or ‘IPRES-enriched’, attained higher TMI than the rest of the patients (Fig. 5A; see Methods). Extending the analysis to the rest of cancer types, we found that prostate, ovarian, lung, and bladder cancers further demonstrated significant difference in TMI between these two predicted groups (p < 0.01 in both platforms; Fig. 5B). As IPRES is predictive of resistance, not response, we further identified 161-gene ‘responder signature’ by applying strict criteria to the DE gene list obtained from the original work (see Methods). Except for melanomas, TMI of all cancers had direct negative correlation with GSVA z-scores of responder signature (Fig. 5C), to different extents, with the most pronounced association seen for lung cancer (Fig. 5D).

**Fig. 5.**
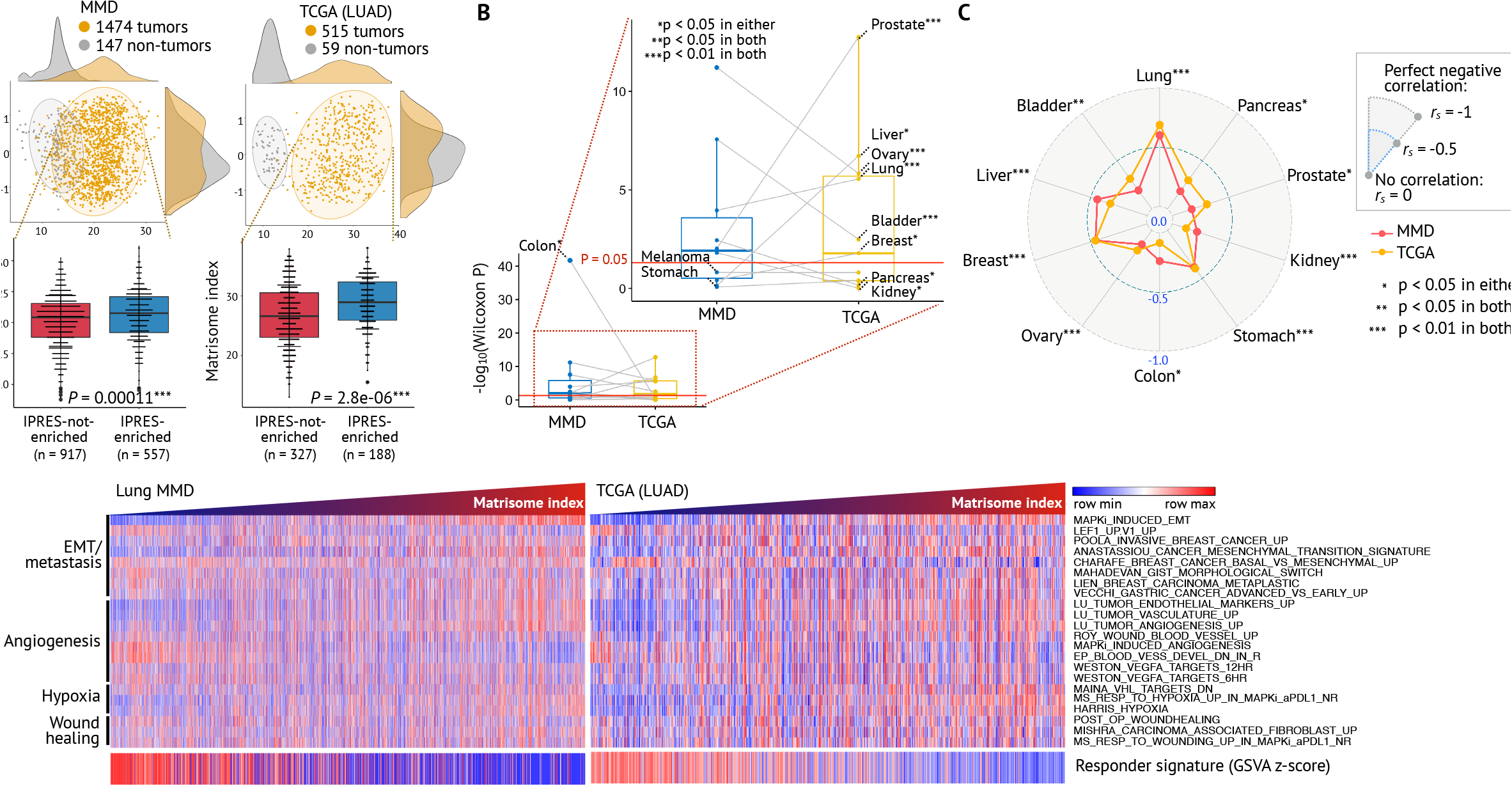
Potential applicability of TMI in predicting immunotherapeutic responsiveness. (A) Statistical significance of the difference between two stratified patient groups according to IPRES (IPRES-enriched: IPRES z-score > 0.35) in lung cancer and (B) the rest of cancer types. (C) Correlation between TMI and GSVA z-scores for responder signature. (D) Heatmaps of GSVA z-scores for IPRES and responder signature in lung cancer. Columns are ordered by increasing TMI.

### Epithelial cancer cells and CAFs are potential cellular contributors of tumor matrisome

We sought to determine the potential cellular source of this common matrisome response observed in bulk tumors. By comparing with previously identified stroma-derived signatures, six ECM genes were reported to be prognostic and differentially expressed in activated stroma, or cancer-associated fibroblasts (CAFs), compared to normal stroma (Table S18) [20, 21]. Many ECM-associated enzymes and other paracrine factors are known to be of fibroblastic origin, leading to altered ECM metabolism and protein expression in tumors [3, 22]. Tumor samples annotated with estimates of both epithelia and stroma components by both pathological and *in silico* means were then analyzed. Tumors with almost no epithelial component expectedly exhibited a wide range of the index, due to stromal cell-derived variation (Fig. 6A). We found an unexpected weak positive correlation between the index and tumor content, indicating tumor cells may also play a role in producing these specific tumor-supporting matrix components (Table S19).

**Fig. 6.**
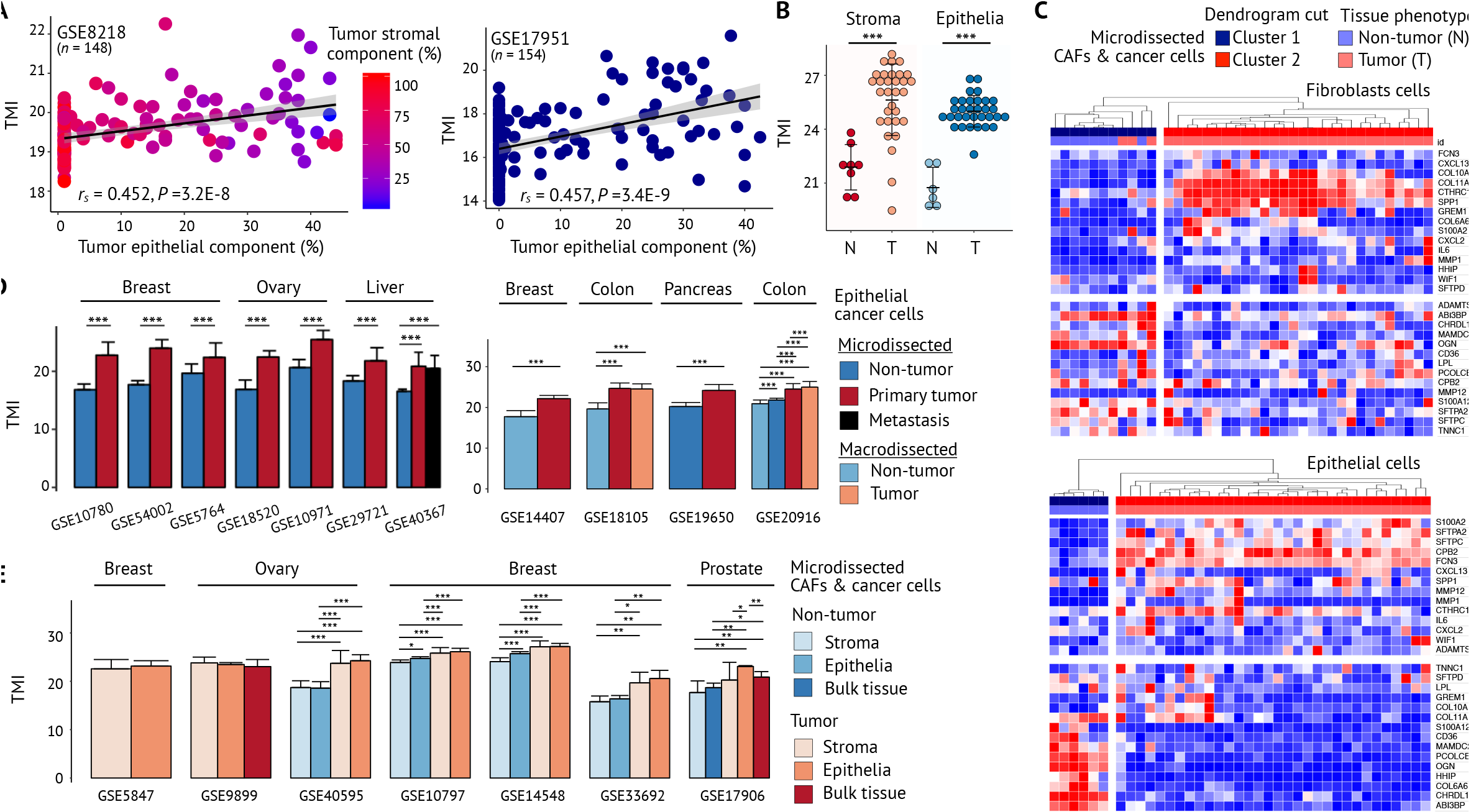
Tumor cells and CAFs as potential cellular contributors of tumor matrisome. (A)Correlation with *in silico* estimates of epithelia and stroma components in prostate cancer. Linear regression lines are drawn (black line) with 95% CI (grey zone). *r*_*s*_ = Spearman’s correlation coefficient; *P* = Spearman’s correlation *P*-value; *n* = number of samples analyzed. (B-C) TMI and heatmaps of laser-microdissected epithelial cells (*n* = 39) and stromal cells (*n* = 38) in high grade serous ovarian cancers from GSE40595. (D-E) Validation using microdissected epithelial and stromal samples in various carcinomas. \*\*\**P* < 0.001, \*\**P* < 0.01, \**P* < 0.05 using the Mann-Whitney U-test.

The initial screening of micro-dissected ovarian tissues revealed significantly higher index in both tumor epithelial cells and stromal CAFs compared to normal ovarian surface epithelial cells and fibroblasts, respectively (Fig. 6B). Heatmaps of these samples demonstrated shared matrisome patterns in both cell types (Fig. 6C). This pattern was also observed in various malignances including breast, ovarian, liver, colorectal, pancreatic, and prostate cancers, showing their ubiquitous alteration at the cellular level (Fig. 6D-E and Table S20). Using human/mouse xenograft models coupled with proteomics, characteristic contribution from both tumor and stromal cells to the production of ECM proteins in tumors of differing metastatic potential was recently reported [23]. The mode by which these cells in tumor matrix produce, degrade, remodel and eventually deregulate tumor matrisome and whether these cell types mutually crosstalk or instruct one another to shape tumor-promoting matrisome remain to be uncovered.

## Discussion

This work is based on cross-platform evaluation of TMI in over 30,000 primary tumors across 11 major cancer types. Prior integrative genome-wide profiling studies are limited to single platforms, interrogating either RNA-seq-based TCGA or microarray-based GEO data. Curating a total of 8,836 patient-derived tumor and tumor-free samples, we generated 11 cancer type-specific MMDs annotated with clinical features and deposited the data at ArrayExpress (see Data Availability). This approach further minimizes biased selection of validation cohorts and allows parallel analyses with other high-throughput data sources - without having to conduct laborious data mining in search for multiple small-scale patient cohorts.

Considering that the index was computed under the assumptions of each gene having concordant regulation as observed in lung cancer, it is noteworthy that tumors with highly dynamic and constantly remodeling ECM could have ubiquitously altered expression of matrisome genes across genetically and phenotypically diverse epithelial tumors. The data may indicate the presence of shared signal transduction pathways activated across these selective tumors, such as recently reported TGFβ or HIF1α/VEGF pathways, in controlling ECM balance or cell-intrinsic mechanism regulating the expression of set of ECM genes contributing to tumor development [23].

Here we provide a comprehensive overview of common matrisome variation in the context of tumor genotypes, molecular and clinical parameters, and immunophenotypes. To the best of our knowledge, this is the first study to suggest potential role of ECM molecules in determining immunosuppressive microenvironment and immune escape mechanisms with cross-platform evaluation of a spectrum of major human malignancies. We found a significant enrichment of tumor-promoting immune infiltrates, IPRES, and multiple immune checkpoints in tumors with high TMI. Heterogeneous infiltration within the same immune cell population was also observed across tumors of varying matrisome indices. For example, high-risk lung tumors according to TMI were significantly enriched for M0 and M1 macrophages, whereas low-risk tumors were negatively enriched for M2 macrophages, suggesting different mechanisms adopted by tumor-associated macrophages (TAMs) to promote tumor progression.

Many of potentially targetable immune checkpoints were enriched in tumors with high TMI, possibly in dulling CD4+ T cell activation, as seen by our CIBERSORT deconvolution analysis. Particularly, we found that B7-H3 emerged as a promising pan-cancer immune target. We expect that such tumors will be more sensitive to immunotherapy due to sufficient immune infiltrates and upregulated expression of immune checkpoints. Tumors with hypermutant phenotypes are regarded as better candidates for immune checkpoint inhibitors across a wide spectrum of cancer types [24–26]. Given positive correlation of TMI with mutational burden, it could thus be postulated that tumors with higher TMI would be more susceptible to immunotherapy. Retrospective analyses using IPRES, however, revealed that high-risk tumors associated more closely with signature predictive of resistance to anti-PD-1 immunotherapy, and vice versa for low-risk tumors. This suggests that high mutational load may not necessarily imply tumor response, as reported in the original study [19]. In particular, lung cancer was found to have the most pronounced correlation effects with IPRES signatures. Given the predictive role of PD-L1 expression for anti-PD-1/PD-L1 therapy in lung cancer [27], enrichment of IPRES in tumors with high TMI may be attributed to insufficient PD-L1 expression in these tumors (Fig. 4C). The findings altogether reinforce the need to adopt a more “holistic” approach to find biomarkers for immunotherapy response that take into consideration tumor, tumor microenvironment and host immunity.

Matrisomal changes during the colorectal and breast cancer multistep carcinogenesis were also implicated. This holds particular promise as no more than a dozen or so “driver” mutations, mostly occurring in tumor suppressors and oncogene, are thought to be potential progression markers of colorectal cancer [28]. Likewise, despite much efforts to characterize ductal carcinoma in situ (DCIS) and invasive ductal carcinoma (IDC), there is currently no standard test accurately identifying tumors likely to progress from preinvasive to invasive carcinomas in breast cancer [29]. Unlike limited information that could be inferred from the binary mutational status, the interpretation of progressively changing TMI on a continuous scale as a progression marker will be clinically pertinent and straightforward.

Since both tumor and CAF are sources of ECM proteins, targeting both cancer cells and CAFs may represent a better strategy to improve antitumor efficacy by remodeling the tumor. Even if actionable targets are found in cancer cells, excessive ECM deposition followed by CAF accumulation presumably build physical barrier and thus prevent efficient drug delivery [30]. Owing to low likelihood of developing clonal selection and therapy resistance, genetic stability of CAFs further makes them promising therapeutic targets [31]. CAF-targeting approach, however, has been challenged for many reasons, such as lack of specific markers and resulting toxicity [32]. The genes that make up this TMI may provide specific targets that are of relevance for this purpose.

Two matrisome-gene-based classifiers and their relevance in clinical outcomes have recently been reported [33, 34]. However, technical feasibility and clinical applicability of these gene panels have not been clearly demonstrated. Their predictive values are further limited to OS, without implication of intrinsic resistance to adjuvant therapy. We showed that the index does not vary greatly with tumor purity without considering other confounding factor (Fig. 6A), suggesting feasibility to the testing. We further demonstrated that the index remains robust in clinical outcomes for multiple surrogate endpoints (OS, DFS, RFS, MFS, PFS, and MO), even in situations where a few genes were missing (table S2), and is an independent predictor of survival. This facilitates easy application and practical development into an affordable multi-gene panel that can be assayed with conventional qPCR in clinical practice. These metrics, nevertheless, will require functional validation in prospective clinical studies and necessitate a standard tissue procession protocols for precise measurement.

## Methods

### Preprocessing of Datasets

All 147 independent public datasets used in this study are summarized in Table S1. Preprocessing methods, number of patient sample, platform assayed, and genes included in the computation of TMI are recorded in Table S2. Raw data of independent studies were RMA-normalized using the *affy* package [35] or preprocessed- or author-defined normalized-data were used as stated in Table S2. Most were assayed with the full 29-gene platform (Affymetrix-GPL570), although some microarray platforms had few missing genes, which are stated in Table S2. As genes without having at least 1 count in at least 20% patients were excluded from preprocessing (described below), different number of ECM genes were included in the computation for the final index for TCGA data. Probes having maximum expression values were collapsed to the genes for subsequent index scoring.

### 12 TCGA Cohorts

Using *TCGA-Assembler* package [36], the Cancer Genome Atlas (TCGA) data of 11 epithelial cancer types were collected, processed, and annotated with clinical parameters (Tables S1 and S2). Due to lack of normal tissue samples, OV and SKCM cohorts, representing ovarian and melanoma cancers respectively, were excluded in DE analyses. *TCGA-Assembler* R package [36] was used to extract normalized RPKM count values. In each dataset, we excluded genes without minimum 1 counts per million (cpm) or RPMK value in less than 20% of total number of samples were excluded using *edgeR* package [37]. These filtered genes were normalized by Trimmed Mean of M-values (TMM) and used for genome-wide DE analyses with *voom* and *lmFit* functions in *limma* package [38]. For survival analyses, clinical information was collected with embedded *DownloadBiospecimenClinicalData* function in *TCGA-Assembler* package [36].

### Generic ECM Signature

We extracted the ranking of 29 ECM genes and visualized their positions in each DE gene list from each cancer-specific TCGA cohort with circular plots generated via the *circlize* package (Table S5) [39]. To further investigate the extent to which ECM genes exhibit greater degree of DE variation among 29 lung-specific ECM genes across all tumors, we derived a “generic” ECM signature based on a weight computed for each ECM gene using the following formula, which was slightly modified from equation that was used to derive generic EMT signature in a previous study [40]:

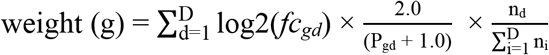

where D is the total number of diseases (D = 6 in this study; prostate, renal, gastric, colorectal, breast, liver, and bladder cancer), *fc*_*gd*_ and *P*_*gd*_ are the fold-change and adjusted *P* value of the ECM gene, g, of disease, *d*, and *n*_*d*_ is the number of patient samples in each TCGA cohort (Table S6). As not all 29 ECM genes were present in the final cancer-specific DE gene list due to preprocessing, each gene was computed with different number of total *n*_*i*_ for the weight. “Generic” ECM signature was then derived from 17 ECM genes having a weighted sum > 3.90 and further visualized for their position in the ranked DE gene list in a circular plot (Fig. 1B). SFTPC gene was not present in the final DE gene list of all TCGA cohorts, except for lung cancer cohort which was not included in the weight computation.

### Tumor Matrisome Index (TMI)

Our 29 ECM gene signature includes collagens (COL11A1, COL10A1, COL6A6), glycoproteins (SPP1, CTHRC1, TNNC1, ABI3BP, PCOLCE2), ECM regulators (MMP12, MMP1, ADAMTS5), ECM-affiliated proteins (GREM1, SFTPC, SFTPA2, SFTPD, FCN3), secreted factors (S140A2, CXCL13, WIF1, CHRDL1, CXCL2, IL6, HHIP, S140A12), proteoglycan (OGN) and other ECM-related components (LPL, CPB2, MAMDC2, CD36) coding genes. Expression profiles of these 29 genes were then extracted from collapsed normalized data and computed for the index, as previously described [7].

### Diagnostic Performance

Complete lists of the patient ID and respective personalized TMI from both tumor and tumor-free samples across microarray and RNA-seq platforms are recorded in tables S7 and S8, respectively. TMI of tumor and normal groups were compared using the Mann-Whitney-Wilcoxon test. To further statistically evaluate the diagnostic accuracy of the TMI in deciding the presence of the disease, we computed the area under the receiver operating characteristic (ROC) curve, sensitivity, and specificity with the best threshold determined by the *pROC* package[41] (Table S9). All ROC curves generated were subjected to binormal smoothing for illustration (Fig. 1D).

### Quantification of TMI Spectrum

We analyzed the Expression Project for Oncology (expO) data provided by the International Genomics Consortium (IGC, USA, www.intgen.com) for ECM spectrum quantification, owing to their procurement of tumor samples under standard conditions. This minimized potential non-biological variations across multiple cancer types, and thus allowed us to perform comparative analysis; data from MMDs and RNA-seq cohorts were excluded for possible batch-effects and different number of genes included in ECM risk scoring, respectively. All patients annotated with prostate, lung, renal, liver, gastric, breast, ovarian, pancreatic, colorectal, and bladder cancer from the expO dataset were included in the spectrum quantification (Fig. 2E and Table S10).

### Patient Stratification

For each preprocessed independent datasets (Table S2), cut-off index was determined using the Cutoff Finder algorithm[42] to stratify each patient cohort into low-and high-risk groups. Statistical summary of 72 datasets used in survival analysis is recorded in Table S11. Kaplan-Meier (KM) survival curves were derived for OS and DSS endpoints (Fig. S1) and other multiple endpoints (Fig. S2) using the *survival* package in R (http://CRAN.R-project.org/package=survival). Multivariate Cox regression analyses were done to adjust confounding factors including age, race, gender, pT, pN, and pM status (Table S12). For all cancer types, patients with available survival data and TMI were all included in the KM analyses.

### Hierarchical Clustering

Heatmaps of samples based on expression profiles using the ECM gene signature were generated with average linkage and Pearson correlation using Morpheus (http://software.broadinstitute.org/morpheus/).

### Molecular Subtypes of Breast Cancer

Of the 147 independent datasets, five datasets (BRCA, GSE20711, GSE21653, GSE19615, and GSE50567) were annotated with either ER, PR, and HER2 status or defined molecular subtypes of breast cancers (normal breast-like, luminal A, luminal B, HER2 positive, and basal-like); tumors harboring positive status of ER and/or negative status of HER2 were classified as luminal cancers while the basal-like tumors were defined as ER-, PR-, and HER2-negative cancers (table S14).

### Total Mutational Burden

We obtained the mutational load of TCGA tumor samples across nine epithelial cancer types from the NCI GDC Data Portal (https://portal.gdc.cancer.gov/projects/) using the same TCGA samples used to generate TMI in our previous analyses. The portal defines the mutational load as the total number of simple somatic mutations. Only tumors harboring at least one mutation were included and log 10 transformed to compute Spearman correlations between TMI and mutational load. To further relate the cut-off of TMI (identified in prior survival analyses) to clinical application, we stratified each TCGA cancer cohort into low-and high-risk groups and compared the mutational load between two groups using the Mann-Whitney-Wilcoxon test (Tables S15 and S16).

### IPRES and GSVA Z-Scores

IPRES was obtained from Broad MSigDB (http://software.broadinstitute.org/gsea/msigdb) and Supplementary Data from the original publication [19]. Using the gene set variation analysis (GSVA), we computed both GSVA score and TMI for any given sample and converted to z-scores. We then obtained the mean z-score for each sample and applied a cutoff of > 0.35, as previously described [19] to define a patient as “IPRES-enriched”. We further defined ‘responder signature’ as 161 highly expressed genes (log FC > 2 and *P* < 0.1) in responders compared to non-responders using initial 693 DE genes identified in the original work [19]. Similar to IPRES, GSVA z-score was computed and correlated with TMI.

### CIBERSORT

The estimated fraction of individual immune cell types was computed using the beta version of CIBERSORT (http://cibersort.standford.edu/). For any given sample, we calculated Spearman correlation between TMI and relative abundance of each immune cell type using our 11 generated MMDs and 11 TCGA cohorts (LUAD, OV, SKCM, BLCA, LIHC, BRCA, COAD, STAD, KIRC, PRAD, and PAAD). As our MMDs of breast, colorectal, and lung cancer exceeded maximum load capacity (500 MB), 1,000 tumors were randomly selected for input data files. We selected LM22 (22 immune cell types) for signature gene file, 100 for permutations, and disabled quantile normalization for all runs.

## Abbreviations

ECM: extra-cellular matrix
NSCLC: non-small-cell lung cancer
PCA: principal component analysis
DE: differential expression
LUAD: lung adenocarcinoma
LUSC: lung squamous carcinoma
PAAD: pancreatic adenocarcinoma
PRAD: prostate adenocarcinoma
KICH: kidney chromophobe
KIRC: kidney renal clear cell carcinoma
KIRP: kidney renal papillary cell carcinoma
STAD: stomach adenocarcinoma
COAD: colon adenocarcinoma
OV: ovarian serous cystadenocarcinoma
BRCA: breast carcinoma
LIHC: liver hepatocellular carcinoma
BLCA: bladder carcinoma
SKCM: skin cutaneous melanoma
TCGA: The Cancer Genome Atlas
CAF: carcinoma-associated fibroblasts
GEO: Gene Expression Omnibus
ExpO: Expression Project for Oncology
ROC: receiver operating characteristic
AUC: area under the ROC curve
OS: overall survival
DSS: disease-specific survival
DFS: disease-free survival
RFS: relapse-free survival
MFS: metastasis-free survival
PFS: progression-free survival
MO: multicentric occurrence
HR: hazard ratio
GSVA: gene set enrichment analysis
GSEA: gene Set Enrichment Analysis
IPRES: innate PD-1 resistance signature
TAM: tumor-associated macrophage.

## Declarations

### Acknowledgements

This work was conceived and carried out at the MechanoBioEngineering laboratory at the Department of Biomedical Engineering, National University of Singapore. We acknowledge support provided by the National Research Foundation, Prime Minister’s Office, Singapore under its Research Centre for Excellence, and Mechanobiology Institute at NUS. We thank both present and former members of the MechanoBioEngineering lab for their support and extend our appreciation to P.Z. and K.J. for proofreading the article. S.B.L. acknowledges scholarship and assistance from NUS Graduate School for Integrative Sciences and Engineering [21].

### Availability of data and materials

Our generated cancer type-specific MMDs are available at ArrayExpress under accession codes E-MTAB-6690 (pancreatic cancer), E-MTAB-6691 (ovarian cancer), E-MTAB-6692 (renal cancer), E-MTAB-6693 (gastric cancer), E-MTAB-6694 (prostate cancer), E-MTAB-6695 (liver cancer), E-MTAB-6696 (bladder cancer), E-MTAB-6697 (melanoma cancer), E-MTAB-6698 (colorectal cancer), E-MTAB-6699 (lung cancer), and E-MTAB-6703 [43].

### Authors’ contributions

S.B.L., S.J.T., W.-T.L. and C.T.L. conceived and designed the study. S.B.L. performed bioinformatics analyses. S.B.L., S.J.T., W.-T.L. and C.T.L. analyzed and interpreted the data. S.B.L., S.J.T., W.-T.L. and C.T.L. reviewed and contributed to the manuscript.

## References

1. Hynes RO. The extracellular matrix: not just pretty fibrils. Science. 2009;326 5957:1216–9. doi:10.1126/science.1176009.

2. Bateman JF, Boot-Handford RP and Lamande SR. Genetic diseases of connective tissues: cellular and extracellular effects of ECM mutations. Nat Rev Genet. 2009;10 3:173–83. doi:10.1038/nrg2520.

3. Lu P, Weaver VM and Werb Z. The extracellular matrix: a dynamic niche in cancer progression. J Cell Biol. 2012;196 4:395–406. doi:10.1083/jcb.201102147.

4. Naba A, Clauser KR, Ding H, Whittaker CA, Carr SA and Hynes RO. The extracellular matrix: Tools and insights for the “omics” era. Matrix biology : journal of the International Society for Matrix Biology. 2016;49:10–24. doi:10.1016/j.matbio.2015.06.003.

5. Hynes RO and Naba A. Overview of the matrisome--an inventory of extracellular matrix constituents and functions. Cold Spring Harbor perspectives in biology. 2012;4 1:a004903. doi:10.1101/cshperspect.a004903.

6. Naba A, Clauser KR, Hoersch S, Liu H, Carr SA and Hynes RO. The matrisome: in silico definition and in vivo characterization by proteomics of normal and tumor extracellular matrices. Mol Cell Proteomics. 2012;11 4:M111 014647. doi:10.1074/mcp.M111.014647.

7. Lim SB, Tan SJ, Lim WT and Lim CT. An extracellular matrix-related prognostic and predictive indicator for early-stage non-small cell lung cancer. Nature communications. 2017;8 1:1734. doi:10.1038/s41467-017-01430-6.

8. Uhlen M, Zhang C, Lee S, Sjostedt E, Fagerberg L, Bidkhori G, et al. A pathology atlas of the human cancer transcriptome. Science. 2017;357 6352 doi:10.1126/science.aan2507.

9. Hoadley KA, Yau C, Wolf DM, Cherniack AD, Tamborero D, Ng S, et al. Multiplatform analysis of 12 cancer types reveals molecular classification within and across tissues of origin. Cell. 2014;158 4:929–44. doi:10.1016/j.cell.2014.06.049.

10. Minn AJ, Gupta GP, Siegel PM, Bos PD, Shu W, Giri DD, et al. Genes that mediate breast cancer metastasis to lung. Nature. 2005;436 7050:518–24. doi:http://www.nature.com/nature/journal/v436/n7050/suppinfo/nature03799S1.html.

11. Gao Q, Wang XY, Zhou J and Fan J. Multiple carcinogenesis contributes to the heterogeneity of HCC. Nature reviews Gastroenterology & hepatology. 2015;12 1:13. doi:10.1038/nrgastro.2014.6-c1.

12. Skrzypczak M, Goryca K,Rubel T, Paziewska A, Mikula M, Jarosz D, et al. Modeling oncogenic signaling in colon tumors by multidirectional analyses of microarray data directed for maximization of analytical reliability. PloS one. 2010;5 10 doi:10.1371/journal.pone.0013091.

13. Toss A and Cristofanilli M. Molecular characterization and targeted therapeutic approaches in breast cancer. Breast cancer research : BCR. 2015;17:60. doi:10.1186/s13058-015-0560-9.

14. Tobin NP, Harrell JC, Lovrot J, Egyhazi Brage S, Frostvik Stolt M, Carlsson L, et al. Molecular subtype and tumor characteristics of breast cancer metastases as assessed by gene expression significantly influence patient post-relapse survival. Ann Oncol. 2015;26 1:81–8. doi:10.1093/annonc/mdu498.

15. Carey LA, Perou CM, Livasy CA, Dressler LG, Cowan D, Conway K, et al. Race, breast cancer subtypes, and survival in the Carolina Breast Cancer Study. Jama. 2006;295 21:2492–502. doi:10.1001/jama.295.21.2492.

16. Tran E, Turcotte S, Gros A, Robbins PF, Lu YC, Dudley ME, et al. Cancer immunotherapy based on mutation-specific CD4+ T cells in a patient with epithelial cancer. Science. 2014;344 6184:641–5. doi:10.1126/science.1251102.

17. Malta TM, Sokolov A, Gentles AJ, Burzykowski T, Poisson L, Weinstein JN, et al. Machine Learning Identifies Stemness Features Associated with Oncogenic Dedifferentiation. Cell. 2018;173 2:338–54 e15. doi:10.1016/j.cell.2018.03.034.

18. Mak MP, Tong P, Diao L, Cardnell RJ, Gibbons DL, William WN, et al. A Patient-Derived, Pan-Cancer EMT Signature Identifies Global Molecular Alterations and Immune Target Enrichment Following Epithelial-to-Mesenchymal Transition. Clin Cancer Res. 2016;22 3:609–20. doi:10.1158/1078-0432.CCR-15-0876.

19. Hugo W, Zaretsky JM, Sun L, Song C, Moreno BH, Hu-Lieskovan S, et al. Genomic and Transcriptomic Features of Response to Anti-PD-1 Therapy in Metastatic Melanoma. Cell. 2016;165 1:35–44. doi:10.1016/j.cell.2016.02.065.

20. Finak G, Bertos N, Pepin F, Sadekova S, Souleimanova M, Zhao H, et al. Stromal gene expression predicts clinical outcome in breast cancer. Nat Med. 2008;14 5:518–27. doi:10.1038/nm1764.

21. Moffitt RA, Marayati R, Flate EL, Volmar KE, Loeza SG, Hoadley KA, et al. Virtual microdissection identifies distinct tumor- and stroma-specific subtypes of pancreatic ductal adenocarcinoma. Nat Genet. 2015;47 10:1168–78. doi:10.1038/ng.3398.

22. Bhowmick NA, Neilson EG and Moses HL. Stromal fibroblasts in cancer initiation and progression. Nature. 2004;432 7015:332–7. doi:10.1038/nature03096.

23. Naba A, Clauser KR, Lamar JM, Carr SA and Hynes RO. Extracellular matrix signatures of human mammary carcinoma identify novel metastasis promoters. eLife. 2014;3:e01308. doi:10.7554/eLife.01308.

24. Rosenberg JE, Hoffman-Censits J, Powles T, van der Heijden MS, Balar AV, Necchi A, et al. Atezolizumab in patients with locally advanced and metastatic urothelial carcinoma who have progressed following treatment with platinum-based chemotherapy: a single-arm, multicentre, phase 2 trial. Lancet. 2016;387 10031:1909–20. doi:10.1016/S0140-6736(16)00561-4.

25. Rizvi NA, Hellmann MD, Snyder A, Kvistborg P, Makarov V, Havel JJ, et al. Cancer immunology. Mutational landscape determines sensitivity to PD-1 blockade in non-small cell lung cancer. Science. 2015;348 6230:124–8. doi:10.1126/science.aaa1348.

26. Snyder A, Makarov V, Merghoub T, Yuan J, Zaretsky JM, Desrichard A, et al. Genetic basis for clinical response to CTLA-4 blockade in melanoma. The New England journal of medicine. 2014;371 23:2189–99. doi:10.1056/NEJMoa1406498.

27. Yu H, Boyle TA, Zhou C, Rimm DL and Hirsch FR. PD-L1 Expression in Lung Cancer. Journal of thoracic oncology : official publication of the International Association for the Study of Lung Cancer. 2016;11 7:964–75. doi:10.1016/j.jtho.2016.04.014.

28. Vogelstein B, Fearon ER, Hamilton SR, Kern SE, Preisinger AC, Leppert M, et al. Genetic alterations during colorectal-tumor development. The New England journal of medicine. 1988;319 9:525–32. doi:10.1056/NEJM198809013190901.

29. Cowell CF, Weigelt B, Sakr RA, Ng CK, Hicks J, King TA, et al. Progression from ductal carcinoma in situ to invasive breast cancer: revisited. Molecular oncology. 2013;7 5:859–69. doi:10.1016/j.molonc.2013.07.005.

30. Chen B, Dai W, Mei D, Liu T, Li S, He B, et al. Comprehensively priming the tumor microenvironment by cancer-associated fibroblast-targeted liposomes for combined therapy with cancer cell-targeted chemotherapeutic drug delivery system. Journal of controlled release : official journal of the Controlled Release Society. 2016;241:68–80. doi:10.1016/j.jconrel.2016.09.014.

31. Qiu W, Hu M, Sridhar A, Opeskin K, Fox S, Shipitsin M, et al. No evidence of clonal somatic genetic alterations in cancer-associated fibroblasts from human breast and ovarian carcinomas. 2008;40:650. doi:10.1038/ng.117 https://http://www.nature.com/articles/ng.117-supplementary-information.

32. Jia D, Liu Z, Deng N, Tan TZ, Huang RY, Taylor-Harding B, et al. A COL11A1-correlated pan-cancer gene signature of activated fibroblasts for the prioritization of therapeutic targets. Cancer Lett. 2016;382 2:203–14. doi:10.1016/j.canlet.2016.09.001.

33. Pearce OMT, Delaine-Smith RM, Maniati E, Nichols S, Wang J, Bohm S, et al. Deconstruction of a Metastatic Tumor Microenvironment Reveals a Common Matrix Response in Human Cancers. Cancer discovery. 2018;8 3:304–19. doi:10.1158/2159-8290.CD-17-0284.

34. Yuzhalin AE, Urbonas T, Silva MA, Muschel RJ and Gordon-Weeks AN. A core matrisome gene signature predicts cancer outcome. Br J Cancer. 2018;118 3:435–40. doi:10.1038/bjc.2017.458.

35. Gautier L, Cope L, Bolstad BM and Irizarry RA. affy--analysis of Affymetrix GeneChip data at the probe level. Bioinformatics. 2004;20 3:307–15. doi:10.1093/bioinformatics/btg405.

36. Zhu Y, Qiu P and Ji Y. TCGA-assembler: open-source software for retrieving and processing TCGA data. Nat Methods. 2014;11 6:599–600. doi:10.1038/nmeth.2956.

37. Robinson MD, McCarthy DJ and Smyth GK. edgeR: a Bioconductor package for differential expression analysis of digital gene expression data. Bioinformatics. 2010;26 1:139–40. doi:10.1093/bioinformatics/btp616.

38. Ritchie ME, Phipson B, Wu D, Hu Y, Law CW, Shi W, et al. limma powers differential expression analyses for RNA-sequencing and microarray studies. Nucleic Acids Res. 2015;43 7:e47. doi:10.1093/nar/gkv007.

39. Gu Z, Gu L, Eils R, Schlesner M and Brors B. circlize Implements and enhances circular visualization in R. Bioinformatics. 2014;30 19:2811–2. doi:10.1093/bioinformatics/btu393.

40. Tan TZ, Miow QH, Miki Y, Noda T, Mori S, Huang RY, et al. Epithelial-mesenchymal transition spectrum quantification and its efficacy in deciphering survival and drug responses of cancer patients. EMBO molecular medicine. 2014;6 10:1279–93. doi:10.15252/emmm.201404208.

41. Robin X, Turck N, Hainard A, Tiberti N, Lisacek F, Sanchez JC, et al. pROC: an open-source package for R and S+ to analyze and compare ROC curves. BMC Bioinformatics. 2011;12:77. doi:10.1186/1471-2105-12-77.

42. Budczies J, Klauschen F, Sinn BV, Gyorffy B, Schmitt WD, Darb-Esfahani S, et al. Cutoff Finder: a comprehensive and straightforward Web application enabling rapid biomarker cutoff optimization. PloS one. 2012;7 12:e51862. doi:10.1371/journal.pone.0051862.

43. Stephens PJ, Tarpey PS, Davies H, Van Loo P, Greenman C, Wedge DC, et al. The landscape of cancer genes and mutational processes in breast cancer. Nature. 2012;486 7403:400–4. doi:10.1038/nature11017.

